# Shortwave-Infrared Line-Scan Confocal Microscope for Deep Tissue Imaging in Intact Organs

**DOI:** 10.1101/2022.12.23.521760

**Authors:** Jakob G. P. Lingg, Thomas S. Bischof, Bernardo A. Arús, Emily D. Cosco, Ellen M. Sletten, Christopher J. Rowlands, Oliver T. Bruns, Andriy Chmyrov

## Abstract

Imaging at wavelengths beyond the visible spectrum enables imaging depths of hundreds of microns in intact tissue, making this attractive for volumetric imaging applications. The development of fluorophores with photoemission beyond 1000nm provide the opportunity to develop novel fluorescence microscopes sensitive to those wavelengths. Here, we present a shortwave-infrared line-scan confocal microscope that is capable of deep imaging of biological specimens, as demonstrated by visualization of labelled glomeruli in a fixed uncleared kidney at depths beyond 400 μm. We also show imaging of brain vasculature labelled with the near-infrared organic dye indocyanine green, the shortwave-infrared organic dye Chrom7, and rare earth-doped nanoparticles, thus encompassing the entire spectrum detectable by a typical shortwave-infrared sensitive InGaAs detector.

## 1. Introduction

One of the current challenges in optical microscopy is volumetric imaging of intact organs at significant depths and high spatiotemporal resolution. Ordinary widefield epifluorescence microscopy, while fast, does not address this challenge as it has inadequate optical sectioning – the ability of an imaging system to reject background fluorescence originating from outside of the focal plane. As a result, the image contrast is reduced, and the detection sensitivity is compromised. While many microscopy techniques [1]–[6] can achieve optical sectioning, they are often restricted by either imaging depth, spatial- or temporal-resolution, or by the geometry of the sample.

One possibility to improve imaging depths in dense tissue is to shift the detection window to longer wavelengths of the electromagnetic spectrum, such as the shortwave-infrared (SWIR, 1000nm to 2500nm) wavelength regime. The SWIR provides several advantages in terms of light-tissue interactions for optical imaging such as reduced scattering, decreased attenuation by blood and pigments, and lower autofluorescence by tissue[7]. Those properties make it an attractive detection window for optical microscopy. The recent developments of SWIR emissive labels[8]–[15] paired with the improvement of SWIR sensitive detectors[16], [17] offer the opportunity to develop novel fluorescence microscopes with optical sectioning capability that are sensitive to wavelengths beyond 1000nm.

SWIR camera-based systems, such as widefield microscopy, have demonstrated *in vivo* imaging depths of up to 800μm in the brain through cranial windows[18]. Further, more advanced widefield techniques requiring image reconstructions have shown millimeter depths through skin and skull [19]. Widefield systems provide high pixel rates of hundreds of gigapixels per second (Gpx/s), however the lack of optical sectioning limits its use to thin or sparsely labelled samples. More advanced methods like light-sheet systems [20], [21] have shown improved optical sectioning by reducing the out-of-focus excitation, nevertheless the off-axis excitation relative to the detection leads to limitations on the possible geometries of samples, making it ineffective for many *in vivo* applications.

In laser-scanning systems, namely confocal and multiphoton[22], the out-of-focus background is suppressed by either the physical rejection of out-of-focus background by pinholes (confocal) or the non-linear signal generation of multiphoton microscopy. Three-photon microscopy has shown imaging depths in the brain beyond 500μm through skull[23], [24] and depths beyond 1mm through cranial windows[25]. However, as a beam is scanned across the sample in two dimensions, the temporal resolution, especially for larger field-of-views, is reduced relative to widefield systems. Moreover, the high photon flux required for three-photon absorption to occur implies lower repetition rates of the laser, typically between 1MHz and 4MHz, compared to the 80MHz repetition rates used in two-photon microscopy. Thus, three-photon microscopy is typically an order of magnitude slower than two-photon microscopy.

SWIR confocal point-scanning systems [16], [17], [26]–[29] have demonstrated imaging depths of 600μm to 1.1mm for various tissues in fluorescence mode and imaging depths of 1.3mm through cranial windows in the brain in reflectance[29]. However, the reported pixel rates of the fluorescence confocal systems have been between 20kpx/s[26] and 260kpx/s[17], [26]. These rates are relatively low compared to the pixel rates achieved by widefield systems. As a result, when scanning, there is a trade-off between spatial and temporal resolution that must be considered.

One possibility to increase temporal resolution while maintaining optical sectioning is line-scan confocal microscopy. As the name implies, in line-scan confocal, a tightly-focused line is scanned across the sample instead of a point; the out-of-focus background is suppressed by a slit or rolling shutter instead of a pinhole[30]–[32]. Line-scan confocal microscopes reduce the scan time per frame as there is no need to scan the line along its axis, which dramatically improves imaging speed at the cost of a slight degradation in the 3D point-spread function.

Here we demonstrate a *de-scanned* shortwave-infrared line-scan confocal microscope (SWIR LSCM) capable of achieving pixel rates of up to 250Mpx/s. The microscope is equipped with an InGaAs line-detector, and is therefore sensitive to shortwave-infrared photoemission of fluorophores. We compare the optical sectioning capabilities of the SWIR LSCM microscope by imaging a liver perfused with fluorescent bacteria on a widefield system and our confocal system. Additionally, we demonstrate the imaging depth of the system by imaging fixed organs, such as the brain and kidneys, labelled with various shortwave-infrared fluorescent probes, namely indocyanine green[10], the chromenylium polymethine dye Chrom7[33] and rare earth-doped nanoparticles[34], at depths beyond 400μm.

## 2. Methods

### 2.1. Optical configuration

The experimental setup of the SWIR LSCM is shown in Fig. 1a. The excitation beam is emitted by a near-infrared fiber-coupled laser (BL976-PAG700, Thorlabs controlled by CLD1015, Thorlabs), this laser beam is collimated by a lens (AC254-075-B-ML, Thorlabs). A cylindrical lens (LJ1640L1-B, Thorlabs) shapes the beam into a line. The beam is reflected upwards by a dichroic mirror (DMLP1000, Thorlabs). After passing a tube lens (TTL200-3P, Thorlabs) and scan lens (SL50-3P, Thorlabs) the beam is scanned by a single-axis silver-coated galvanometer (GVS001, Thorlabs). The beam passes a second pair of scan and tube lenses before forming the illumination line in the back-focal plane the objective (25X XLSLPLN25XGMP NA 1.0, Olympus; with 4:1 glycerol (3783.1, Roth) to water (Millipore) as immersion medium). The fluorescence signal is epi-collected by the high NA objective and de-scanned by the galvanometer. The fluorescence signal passes the dichroic mirror and is spectrally filtered by an emission filter (FELH1000, Thorlabs). An objective lens (4X N4X-PF NA 0.13, Nikon) forms the fluorescence image on the high-speed line-camera (Manx 1024 R, Xenics) attached to a manual x,z-stage (DTS50/M and PT1/M, Thorlabs). The use of the line-detector removes the need for a slit or a rolling shutter to achieve optical sectioning. We decided to use an optical configuration that does not lead to diffraction limited imaging and does not make use of the full detector width. The reason is that the larger field of view paired with lower resolution allows for more fluorescence signal to be collected and enables faster imaging on the high speed InGaAs line detector.

**Fig. 1:**
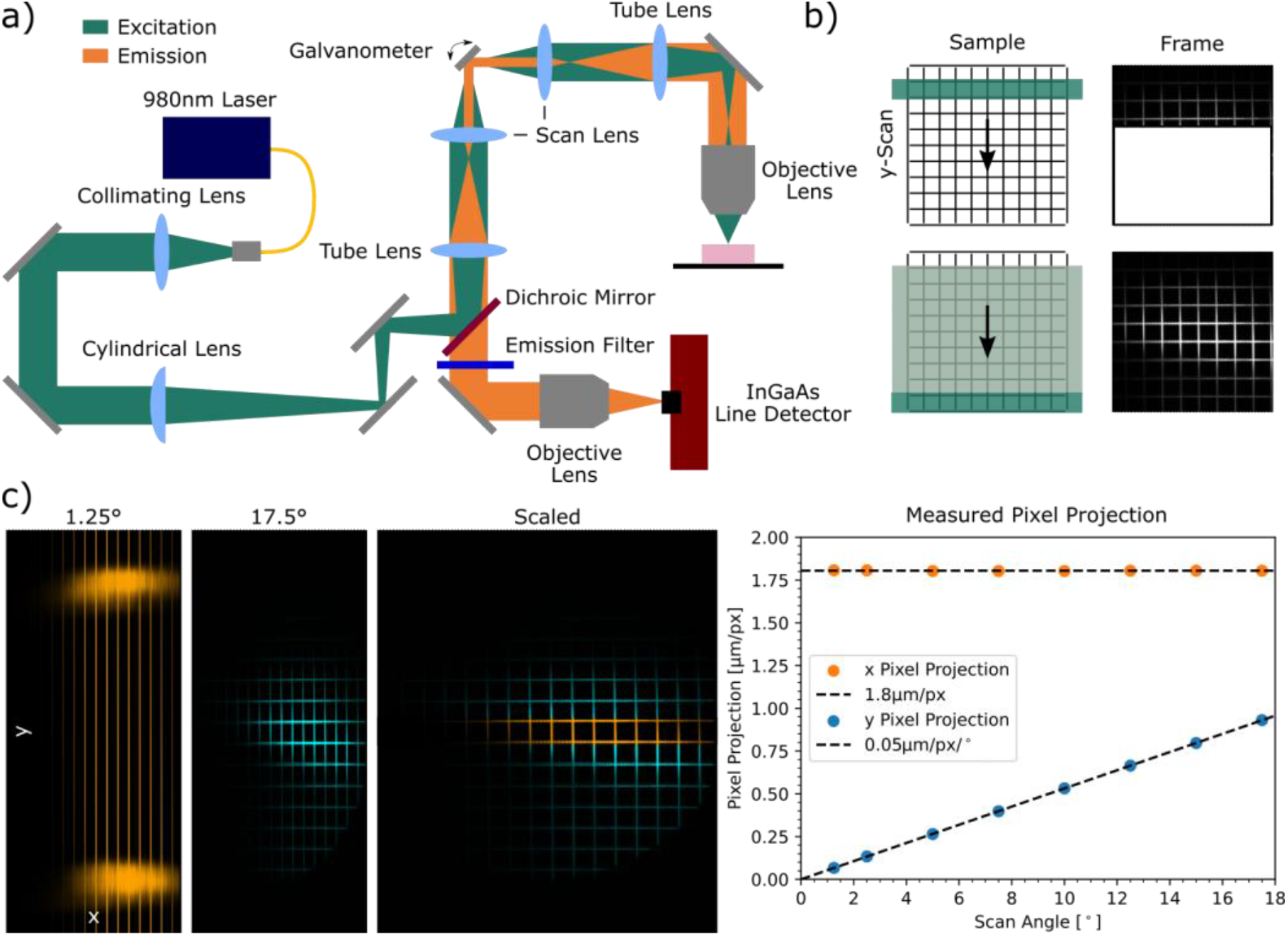
SWIR LSCM setup and working principle. a) Schematic of SWIR line-scan confocal microscope. b) Explanation of the scan mechanism. A line is scanned from top to bottom (y-axis), the line-detector acquires lines and stitches lines to form the final frame. c) 50μm grid imaged at different scan angles (1.25°, 17.5°), including the overlay of the scaled frames. Plot shows the measured pixel-projection per scan angle for the x- and y-axis.

The camera receives two digital signals: the first signal to initialize a frame, followed by signals to trigger the acquisition of single lines. The lines are collected and stitched along the y-axis forming a full frame; the scanning mechanism is shown in Fig. 1b. Frames are acquired for both the forward and backward scan directions to maximize the frame rate.

To stabilize the temperature of the thermoelectric cooled sensor, we attached a heat sink to the detector and positioned a cooling fan (H17543-001, Intel) close to the detector to be able to image consistently at -10°C detector temperature, resulting in a detector sensitivity range of 900nm to 1650nm. A digital acquisition (DAQ) card (PCIe-6351, National Instruments) generated the electrical signals used to drive the galvanometer (triangular wave) and trigger the acquisitions of the camera.

The z-scanning by the microscope is performed by moving the objective mounted on a fixed-single-objective nosepiece (CSN100, Thorlabs) with a motorized focusing module (ZFM2020, Thorlabs). The electronic components (galvanometer, camera, x,y-stage, z-stage) and the acquisition of stacks is controlled by self-written custom software with a graphical user-interface (Qt C++). The microscope control code is available at GitLab (https://gitlab.com/brunslab/swir-line-scan-confocal). Data exploration was done using ImageJ. Data processing, analysis and plotting was performed with Python using the code, which is available at https://gitlab.com/brunslab/manuscript-swir-line-scan-confocal.

The acquired stacks were dark corrected and the frames were divided by the normalized median of a substack of the stack to reduce the detector contributions. Two columns of pixels were removed for the nanoparticle frames shown because the pixels were blinking at the high exposure times applied.

#### ICG, Nanoparticle, and Chrom7 Image Acquisition

For the ICG, nanoparticle, and Chrom7 data acquisition, we swapped the objective lens positioned at the detector with a 2.5X lens (2.5X PE IR Plan, Seiwa). Further, we used an achromat with 50mm focal length (ACL254-050-B-ML, Thorlabs) to collimate the beam.

For the nanoparticle image acquisition, we used an immersion oil (n=1.48 Type FF, Cargille) instead of a glycerol solution and placed a 1350nm long-pass filter (FELH1350, Thorlabs) into the detection path.

Solely for the ICG images, we used a 785nm fiber-coupled laser (S4FC785, Thorlabs with P3-780A-FC-2, Thorlabs) spectrally cleaned by a short-pass filter (FESH0800, Thorlabs), replaced the two tube lenses (TTL200-2P, Thorlabs), used a different dichroic (DMLP805R, Thorlabs), and used an 850nm long-pass filter (FELH0850, Thorlabs) in the detection path. Additionally, we changed one of the scan lenses (SL50-2P, Thorlabs).

### 2.2. Pixel Projection Characterization

As the scan range of the galvanometer and the number of lines to be collected per frame can be varied, the resulting images need to be scaled accordingly. Fig. 1c displays the frames acquired in reflectance of the same grid (50μm R1L3S3P, Thorlabs) before and after scaling for different scan angles for the same number of lines per frame (1024 lines). The pixel projection per scan angle of the galvanometer is determined by imaging the grid using a variety of scan angles in reflection. The images are background corrected to remove the signal attained from internal reflections. For each background corrected image, the sum along the y-axis and the x-axis is taken. The sums are plotted and the peaks in the image, corresponding to grid lines, are found. The distance between the peaks in pixels corresponds to the known distance of the grid lines, in this case 50μm. Thus, we find the pixel projection per frame in the x- and y-axis. We fit a constant to the pixel projection in the x-dimension per scan angle and a linear function to the y-dimension pixel projection per scan angle. The x-axis pixel projection corresponds to the theoretical pixel projection of 1.8 μm/px. The y-axis pixel projection per scan angle is 0.05 μm/px/°. With those coefficients the acquired images can be scaled for any scan angle.

### 2.3. Bacteria Resolution Target

Due to the lack of commercially available point-sources (fluorescent beads, etc.) with photoemission beyond 1000nm, we estimated the resolution of the microscope by measuring SWIR fluorescent bacteria (*Blastochloris viridis*, DSM no. 133, DSMZ)[35] embedded in agarose. Fig. 2a shows the absorption and emission spectra of the bacteria with peaks at 1012nm and 1044nm, respectively. From a widefield image (IX83, 100X UPLXAPO100X NA 1.45, Olympus; Imaging Source DFK MKU226-10×22) of the crystal violet-stained bacteria (Fig. 2b), we estimate the length of a single bacterium to be between 1.5μm and 2.5μm, the width is estimated to be between 0.5μm and 0.8μm.

**Fig. 2:**
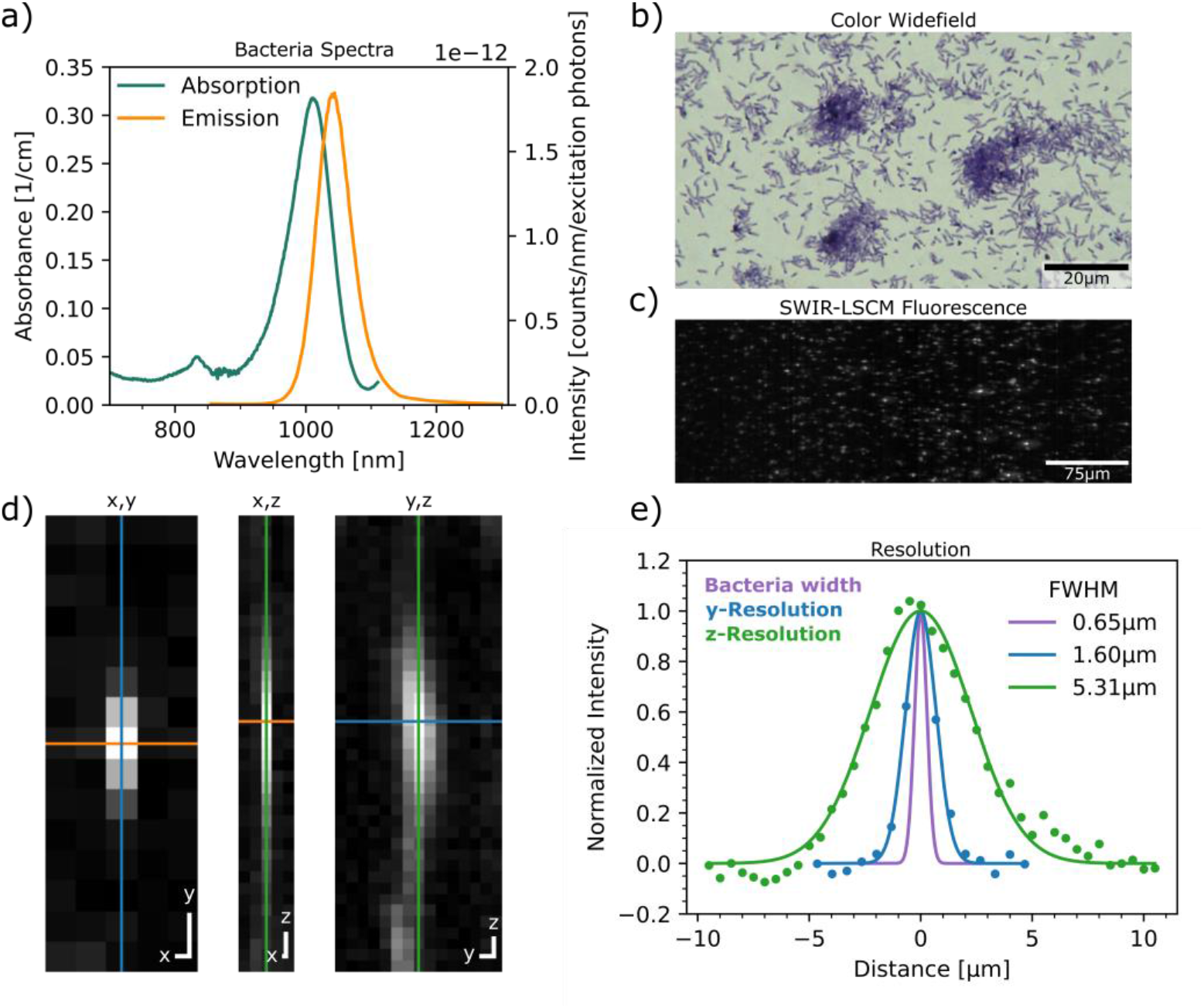
Resolution estimation using fluorescent bacteria. a) Absorption and emission properties of the bacteria. b) *Blastochloris viridis* stained with crystal violet (IX83 with 100X UPLXAPO100X NA 1.45, Olympus; Imaging Source DFK MKU226-10×22). c) SWIR LSCM acquired maximum intensity projection of 50μm of bacteria embedded in agarose (excitation wavelength: 980nm). d) Unscaled image planes of single bacterium imaged with SWIR LSCM as resolution estimate (1μm scale bar). e) Resolution estimate plot including the bacterium size as the width of a stained bacterium (purple line), resolution measurement in y-dimension (confocal direction, blue line), and the resolution measurement in z-dimension (green line).

A vial with bacteria in medium (Rhodospirillaceae medium no. 27, DSMZ) was centrifuged, the excess medium removed, and Dulbecco’s Phosphate Buffered Saline DPBS (14190-094, Gibco) was added to the bacteria. The bacteria solution was vortexed for several minutes until no bacteria clusters were visible in the solution. Bacteria solution was mixed with 1% agarose solution in a 35mm glass bottom dish (81218-200, Ibidi). Once the agarose-bacteria mixture solidified, the glass bottom dish was attached to a custom 3D-printed container which then was filled with the immersion medium (4:1 glycerol (3783.1, Roth) to water (Millipore)).

A stack with axial step size of 0.5μm was acquired. The stack was dark corrected, before being normalized for bleaching by dividing the frames with the average frame value of the section; this assumes a homogenous and isotropic distribution of bacteria (a maximum intensity projection of the first 50μm depth is shown in Fig. 2c). A rolling ball background correction with radius 10px was then applied. A Gaussian curve was fitted through the values of the line-profiles passing through the brightest spot of the bacteria sub-stack (Fig. 2d). The full-width half-maximum of the y-axis resolution (confocal direction) was measured to be 1.6μm, the z-resolution to be 5.3μm (Fig. 2e).

### 2.4. Absorption and Emission Spectra

#### Absorption Spectrum of Bacteria

The absorption spectrum of the fluorescent bacteria (Fig. 2a) was measured with a UV/VIS/NIR spectrophotometer (Lambda1050+, PerkinElmer) equipped with a 150mm WB-InGaAs (wideband) integration sphere (L6020364, PerkinElmer). We measured the spectrum of the bacteria in medium in a plastic cuvette (Hach 2629500, 10mm path length, 1.5mL); the acquired spectrum was corrected by subtracting the spectrum of the pure medium.

#### Emission Spectra of Bacteria, ICG, Nanoparticles, and Chrom7

We measured the emission spectra of the samples using a liquid nitrogen cooled InGaAs line-detector (Pylon-IR 1024-1.7, Princeton Instruments) and a Silicon detector (PIXIS: 400BR, Princeton Instruments) attached to a Czerny-Turner spectrograph (HRS-300, Princeton Instruments). ICG, *Blastochloris viridis*, and the rare earth-doped nanoparticles were excited with 635nm, 785nm, and 980nm wavelength, respectively (SuperK-Extreme, NKT Photonics filtered with SR-Extended-8HP LLTF Contrast, NKT Photonics). Additionally, the excitation light was spectrally filtered by short-pass filters (ICG: FESH0650, Thorlabs; *Blastochloris viridis*: FESH0800, Thorlabs; rare earth-doped nanoparticles: FESH1000, Thorlabs) and focused on to the mounted cuvette (Hach 2629500, 10mm path length, 1.5mL) by an achromat (AC254-200-AB-ML, Thorlabs). The photoemission was collected using a pair of silver-coated off-axis parabolic mirrors (MPD269-P01 and MPD249H-P01, Thorlabs) that focus the light onto a multimode fiber (400μm, 0.39 NA, M28L01, Thorlabs) that enters the spectrograph. Long-pass filters (ICG: FELH0650, Thorlabs; *Blastochloris viridis*: FELH0850, Thorlabs; rare earth-doped nanoparticles: FELH1150, Thorlabs) were placed in front of the fiber to suppress the excitation light. Acquired spectra were background corrected and intensity corrected using a calibration lamp (SLS201L/M, Thorlabs), considering the quantum efficiency of the detector and wavelength dependent transmission losses of the system components. The bacteria spectrum was wavelength binned and normalized for the excitation power (ICG: 0.3mW; *Blastochloris viridis*: 4mW; rare earth-doped nanoparticles: 3.9mW) at the sample, resulting in a spectrum in units of counts per wavelength per number of excitation photons. The spectral data of Chrom7 was obtained from the authors of its publication [33].

### 2.5. Animal preparation

#### Bacteria Perfusion

A mouse (Athymic Nude (Crl:NU(NCr)-Foxn1nu), female, 7 weeks, Charles River) was anesthetized with an i.p. Ketamin Xylazin injection. The blood was flushed out by perfusing the mouse with heparin in PBS (20U/ml, 375095-100KU, Merck) through the heart using a continuous flow syringe pump (4ml/min, 4000-X, Chemyx). Afterwards, 7.5ml *Blastochloris viridis* in PBS (14190-094, Gibco) was flushed through the vasculature using the syringe pump. Organs were extracted and fixed in Paraformaldehyde solution 4% in PBS (SC-281692, Santa Cruz Biotechnology).

#### ICG Injection

A nude mouse (Crl:NU(NCr)-Foxn1nu, female, 6 weeks, Charles River) was injected intravenously with 200μl of 0.05mg/ml ICG (7695.3, Roth) in water (L0970-500, Biowest) and immediately sacrificed. The brain was extracted and fixed in Paraformaldehyde solution 4% in PBS (SC-281692, Santa Cruz Biotechnology).

#### Nanoparticle Injection

A nude mouse (Crl:NU(NCr)-Foxn1nu, female, 6 weeks, Charles River) was injected intravenously with 200μl of 20mg/ml rare earth-doped nanoparticles (NBDY-0029A, NIRmidas) in PBS and immediately sacrificed. The brain was extracted and fixed in Paraformaldehyde solution 4% in PBS (SC-281692, Santa Cruz Biotechnology).

#### Chrom7 Injection

A nude mouse (Crl:NU(NCr)-Foxn1nu, female, 8 weeks, Charles River) was injected intravenously with 200μl of 0.64mg/ml Chrom7 formulated into water-soluble poly(ethylene) glycol-phospholipid micelles in saline [33], and immediately euthanized. The brain, kidney, and liver were extracted and fixed in paraformaldehyde 4% in PBS (SC-281692, Santa Cruz Biotechnology).

### 2.6. Widefield imaging

We used an inverted widefield microscope (IX83, Olympus) with an InGaAs detector (Nirvana HS, Teledyne or Goldeye-034, Allied Vision) to make the comparison between the SWIR LSCM and the widefield system (Fig. 3). The microscope was used to image the cryo-sliced sections of the kidney (Supplementary Figure 4) in fluorescence and brightfield configuration (absorption contrast). Excitation light of a near-infrared laser (980nm, LU0980D350-U30 AF, Lumics) was coupled via a 30:70 beamsplitter (BS020, Thorlabs) to a light guide into the microscope body. The filter cube used consists of a 1000nm short-pass filter (FESH1000, Thorlabs), dichroic mirror (DMLP1000, Thorlabs), and a long-pass filter (FELH1050, Thorlabs) in the detection path.

**Fig. 3:**
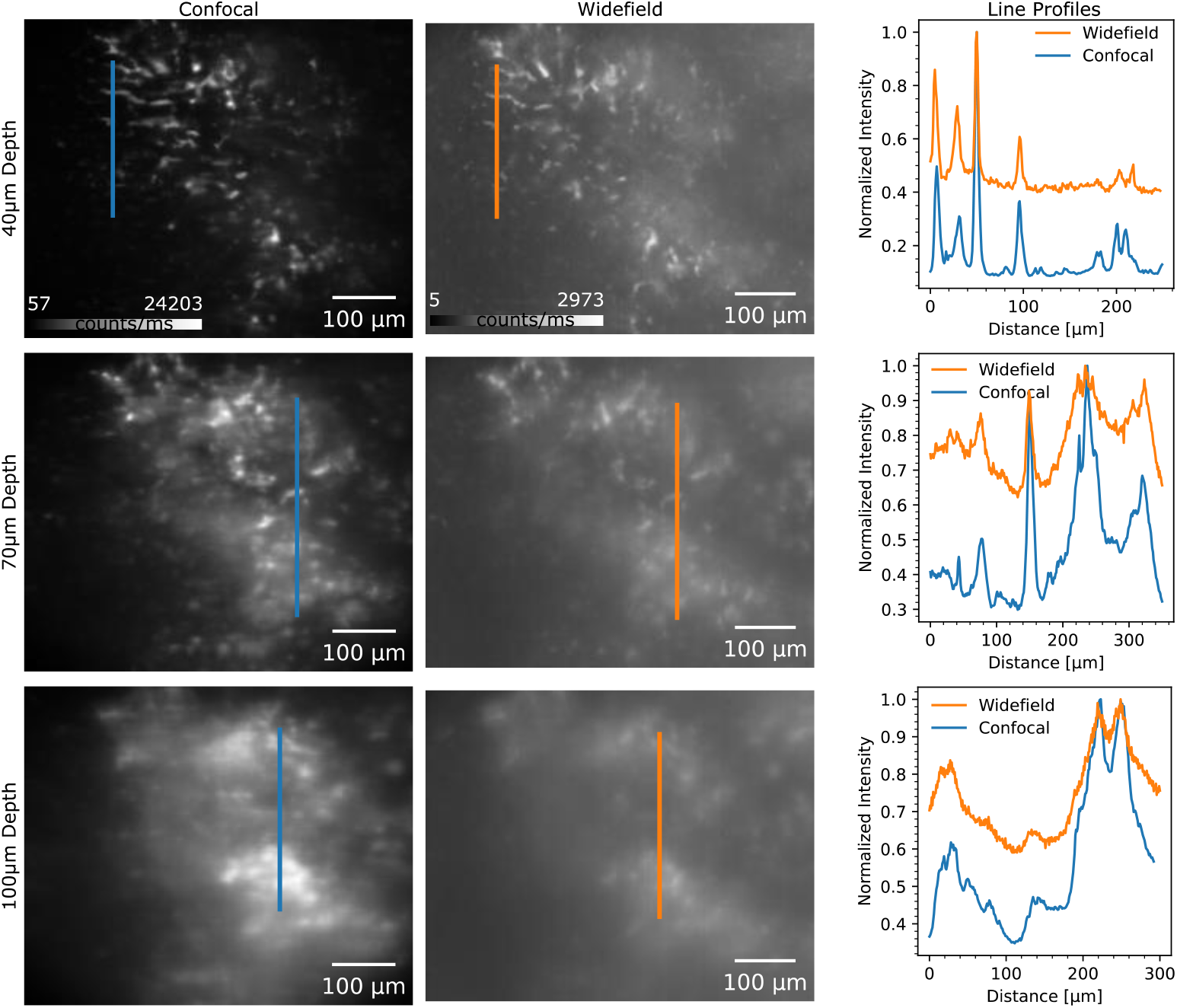
Optical sectioning capabilities of SWIR LSCM. Left and center columns: Images of mouse liver perfused with bacteria at different depths taken with both the SWIR LSCM and the widefield system. The colorbar indicates the signal in counts per exposure time; the line exposure time for the SWIR LSCM was 2.2ms (excitation wavelength: 980nm, frame height: 1024 lines, galvo frequency: 0.2Hz, detector width used: 500px, pixel rate: 205kpx/s, detector gain: 20×) and the widefield exposure time was set to 20ms on the NIRvana HS, Teledyne. Right column: Intensity-normalized line profiles as shown in the SWIR LSCM and widefield images.

## 3. Results

### 3.1. Comparison to Widefield Microscopy

We evaluated the optical sectioning capabilities of the SWIR LSCM by imaging the same structure in a fixed mouse liver labelled with fluorescent bacteria on a widefield system and the confocal system (Fig. 3). We embedded the liver in agarose in a 35mm glass bottom dish (81218-200, Ibidi) to simplify the transition and correlation between the inverted widefield system (20X UPLXAPO20X NA 0.8, Olympus) and the confocal system. A narrow tape was adhered on top of the cover glass to allow positional alignment of the sample in both systems via brightfield imaging on the widefield system and reflectance imaging on the confocal.

We matched the pixel projection of the widefield system (∼1μm/px) by adjusting the scan angle of the confocal system. Both stacks were background and flat corrected. The acquired frames had sufficient signal to noise, therefore no additional rolling ball background subtraction was applied. From the images alone, here normalized to minimum and maximum values, the contribution of the out-of-focus background is noticeably stronger in the widefield images compared to the confocal images. Vertical line-profiles normalized to the maximum were plotted, indicating that the contrast in the confocal images is approximately a factor of three greater than in the images taken with the widefield system.

### 3.2. Kidney Imaging with SWIR LSCM

To further understand the capabilities of the SWIR LSCM, we imaged a fixed mouse kidney whose vasculature was labelled with the shortwave-infrared fluorescent chromenylium polymethine dye Chrom7[33]. Chrom7 has an emission peak at around 1000nm (Supplementary Figure 1). The acquired stack, shown in Fig. 4 displays the vasculature of the surface layers of the kidney, and beyond 240μm of imaging depth, the glomeruli, the structures in which blood is filtered, become visible. We were able to image glomeruli at depths past 400μm within the fixed kidney; the full stack is displayed in Supplementary Figure 2. The appearance of glomeruli in the images at varying depths again highlights the instrument’s optical sectioning capabilities. The diameter size of the glomeruli between 40μm to 100μm and the position of the glomeruli at depths beyond 200μm from the kidney surface corresponds to known literature values[36]. We color-coded the stack slices according to the depth[37] and performed a maximum intensity projection (MIP) of the colored stacks. This allows us to see the contributions of labelled vessels close to the surface of the kidney and the glomeruli deep within the kidney.

**Fig. 4:**
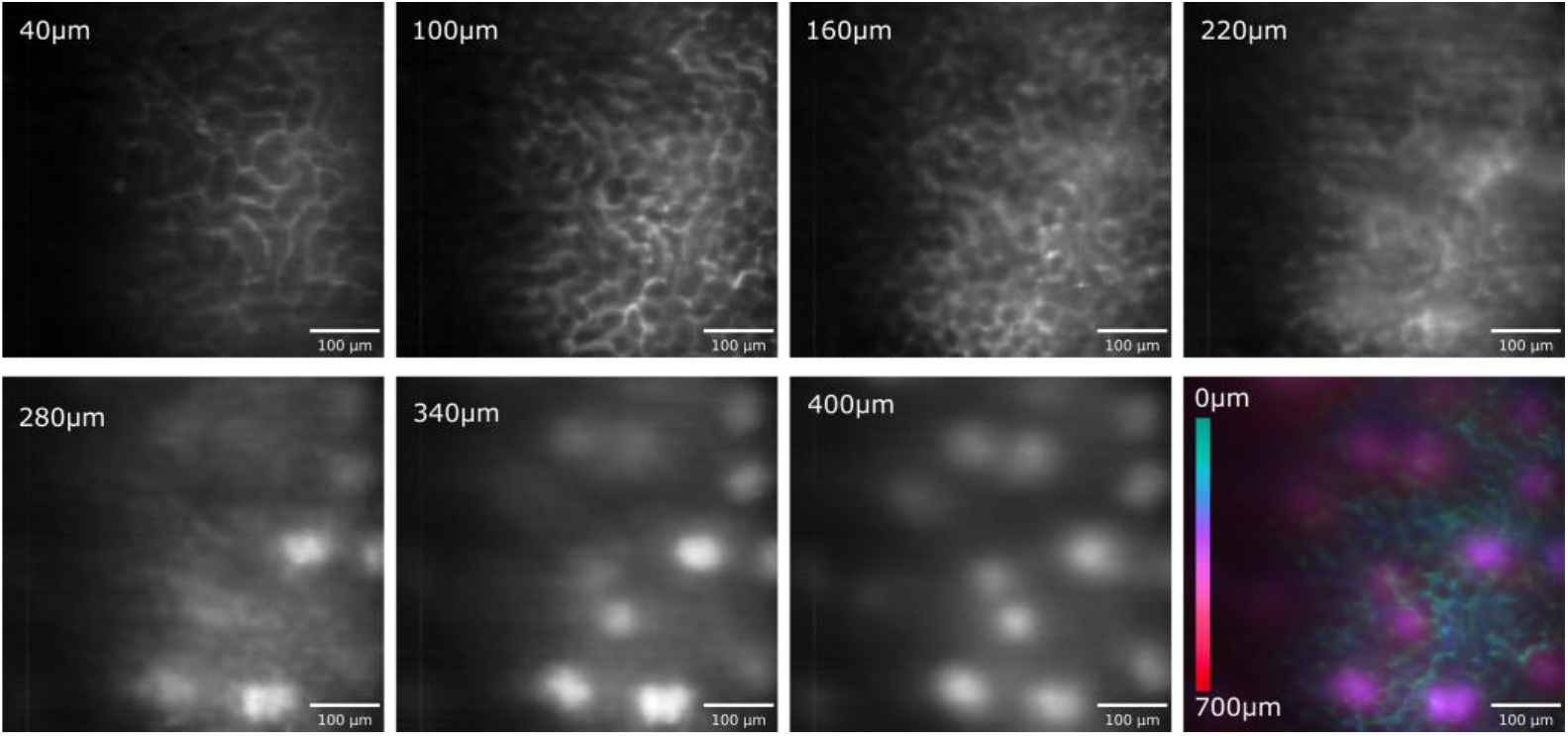
Kidney imaging using SWIR LSCM. Sections of intact fixed kidney labelled with Chrom7 imaged with SWIR LSCM; the line exposure was set to 11ms (excitation wavelength: 980nm, frame heigth: 512 lines, galvo frequency: 0.08Hz, detector width used: 730px, pixel rate: 60kpx/s, detector gain: 20×). Glomeruli are visible in frames past 240μm depth.

In addition to imaging a kidney stained with Chrom7, we imaged a mouse kidney perfused with the fluorescent bacteria *Blastochloris viridis*, which showed an accumulation of the bacteria in the glomeruli (Supplementary Figure 3). We verified the labelling of glomeruli with bacteria by imaging cryo-sliced sections of the kidney in fluorescence on the widefield system (using an Olympus 4X UPLAXAPO4X NA 0.16 objective) [36] and performing hematoxylin and eosin (H&E) staining of the same section (Supplementary Figure 4).

### 3.3. Brain Imaging with SWIR LSCM

We further demonstrate the capabilities of the SWIR LSCM by imaging mouse brain vasculature labelled with the clinically-approved near-infrared dye indocyanine green (ICG), shown in Fig. 5a. Upon near-infrared excitation, ICG exhibits strong photoemission well beyond its emission peak (about 830nm in Methanol, see Supplementary Figure 1). Prior work has demonstrated that it is possible to image ICG fluorescence in the spectral range 900nm to 1600nm using InGaAs area detectors[10]. Here, we employ the InGaAs line detector in our SWIR LSCM, which offers sensitivity from 900nm to 1650nm, allowing us to capture images of ICG. The acquired image stack shows a dense vascular structure close to the brain surface, paired with larger vessels. Deeper within the brain, the larger vessels disappear, and capillaries become visible. We were able to reach an imaging depth of 400μm before losing imaging contrast, as seen in the full stack in Supplementary Figure 5.

**Fig. 5:**
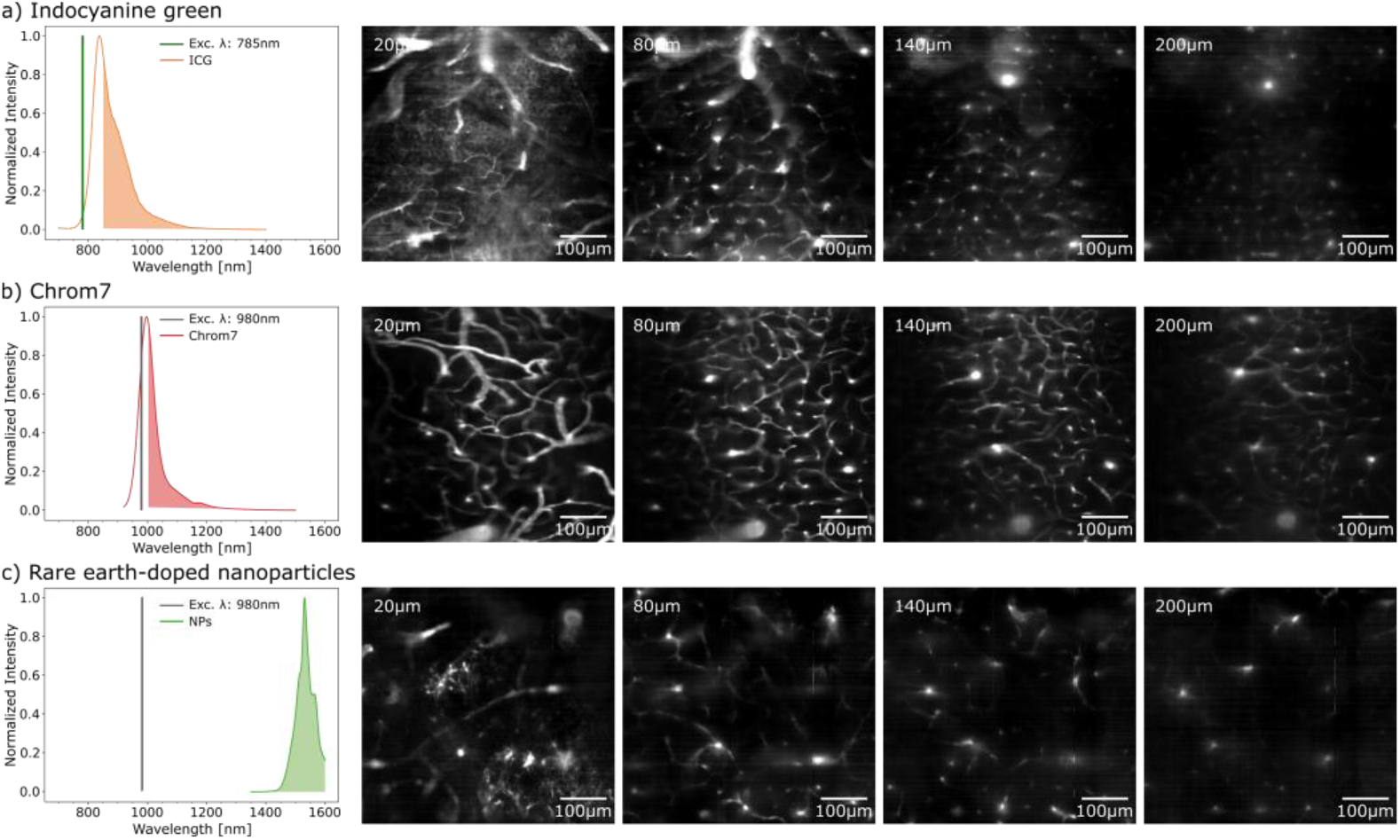
Brain vasculature imaging using SWIR LSCM with various labels including corresponding emission spectra. a) Sections at various depths of fixed mouse brain with ICG-labelled vasculature imaged with 8.8ms line exposure time (excitation wavelength: 785nm, frame height: 512 lines, galvo frequency: 0.1Hz, detector width used: 730px, pixel rate: 75kpx/s, detector gain: 100×, emission filter: FELH0850, Thorlabs). b) Sections of fixed mouse brain labelled with the organic dye Chrom7 imaged with 8.8ms line exposure time (excitation wavelength: 980nm, frame height: 512 lines, galvo frequency: 0.1Hz, detector width used: 730px, pixel rate: 75kpx/s, detector gain: 100×, emission filter: FELH1000, Thorlabs). c) Sections of rare-earth doped nanoparticle labelled mouse brain imaged with 100ms line exposure time (excitation wavelength: 980nm, frame height: 512 lines, galvo frequency: 0.008Hz, detector width used: 730px, pixel rate: 6kpx/s, detector gain: 100×, emission filter: FELH1350, Thorlabs).

As a substitute to ICG, we utilized the shortwave-infrared organic dye Chrom7 to label brain vasculature (Fig. 5b). Our realized imaging depth was around 400μm before losing imaging contrast, and we could not distinguish capillaries anymore. The full stack is available in Supplementary Figure 6. Further, it appears that the mononuclear phagocytic system takes up Chrom7, which is present in organs that are rich in macrophages, such as the liver. The maximum intensity projection of a stack of liver images displayed in Supplementary Figure 7b reveals cellular labeling with Chrom7.

Finally, we imaged a fixed mouse brain labelled with rare earth-doped nanoparticles (NBDY-0029A, NIRmidas)[34] (Fig. 5c) with an emission peak at 1550nm wavelength (Supplementary Figure 1). While the nanoparticles are not prone to bleaching, their inferior brightness, when compared to the acquired ICG or Chrom7 stacks, leads to a loss of detected fluorescence signal, eventually dropping below the noise floor of the InGaAs line-detector when imaging deeper structures. Nonetheless, we were able to image at depths up to 400μm (Supplementary Figure 8).

## 4. Discussion

We developed a microscope that can image hundreds of microns deep into fixed uncleared tissue, which opens many new opportunities in terms of deep tissue imaging in intact dense organs. The speed of our current system is limited by the comparatively-high noise floor of the InGaAs detector relative to the noise levels of standard silicon detectors, and not yet by the scanning speed of the galvanometer or the line-rate of the camera. SWIR detectors undergo rapid developments, resulting in notable reductions in noise levels and, consequently, improved detection sensitivity. Another possibility to overcome the mentioned challenge is to use labels with increased SWIR brightness, such as aggregation-induced emission (AIE) dots[38]. The high brightness of AIE dots was demonstrated to be vital for enabling biomedical applications such as mapping physiological and pathological functions of cortical vasculature and its response to stroke and thrombosis in animals[39]. Organic dyes like ICG, while having lower brightness per emitter, have the significant advantage of being clinically approved, significantly reducing the regulatory burden for potential medical applications in humans. The parallelized acquisition allows to trade slower scanning speed for longer acquisition times, thus enabling the use of organic fluorophores.

Versus multiphoton microscopy, the SWIR LSCM provides more penetration depth compared to two-photon measurements in a fixed kidney of up to 200μm (see Supplementary Figure 9), which are in line with reported two-photon measurements (∼200μm) in kidneys[40]. The linear excitation of the line-scan confocal system leads to larger contributions of out-of-focus excitation, where the laser excitation is sufficient to excite the fluorophores in superficial structures of the tissue, even though it is not focused on this position, reducing the image contrast. Such loss of contrast is also forming a limit for achievable imaging depths in multiphoton microscopy [41]. In addition, multiphoton processes suffer from loss of efficiency due to pulse broadening by dispersion, which does not affect single photon excitation in confocal microscopy. The imaging speed of the multiphoton systems, as any point-scanning technique, is limited by the scanning speed of the beam scanners and emission rates from the labels. Thus, the field-of-view must be restricted to be able to image at high temporal resolution. We overcome this issue by parallelizing the acquisition on a line-detector. Generally, our microscope is more cost-effective and easier to implement compared to multiphoton microscopes. Further, line-scan confocal achieves superior optical sectioning capabilities compared to multi-spot confocal and spinning disk confocal systems[42].

SWIR-LCSM is a new technique and there is potential for improvement. Adding more excitation wavelengths to enable the use of multiple fluorophores via excitation multiplexing in the SWIR[11] and the addition of a reflectance channel would improve the usability of the instrument. The installation of the reflectance channel would require the adding of polarizing optics to suppress the contribution of internal reflectance arising from the optics[29], potentially leading to transmission losses. To take better advantage of the InGaAs line-detector’s width and to be able to increase the pixel projection [μm/px], a different imaging lens, such as one with a 300mm focal length, could be utilized. This would result in a pixel projection of 0.67μm/px. However, given our current signal limitations, we have not decreased the pixel projection in our system any further. As the SWIR spectral range is not commonly used in microscopy, additional attention should be paid to the transmission properties of all optical components.

In conclusion, SWIR LSCM provides an imaging speed increase of at least a factor of four over point-scanning systems. We achieved pixel rates of 205kpx/s with a detector gain setting of 20× and only using half of the detector width when imaging fluorescent bacteria. Using the same fluorescent label, we can improve the imaging speed further by increasing the detector gain to 100×, leading to pixel rates of 1Mpx/s while tolerating increased noise levels. Comparable SWIR confocal systems [16], [17], [26] report pixel rates between 20kpx/s and 260kpx/s in fluorescence. Thus, we believe that with the current state of SWIR photoemissive labels and SWIR sensitive detectors, the line-scan imaging configuration is the most viable option to perform optical sectioning in the SWIR with potential dynamic imaging applications in mind.

## Acknowledgments

We thank Hannes Rolbieski for the support with the animal handling, mouse perfusion, and injection. Further, we thank David Kamel for the support with the bacteria culture.

## Data availability

Datasets, including all raw and processed imaging data generated in this work, will be made available at the BioImage Archive with the publication.

## Code availability

Source code for the microscope control is available at https://gitlab.com/brunslab/swir-line-scan-confocal. The processing and analysis scripts are available at https://gitlab.com/brunslab/swir-line-scan-confocal.

## References

1. J.-A. Conchello and J. W. Lichtman, ‘Optical sectioning microscopy’, Nat Methods, vol. 2, no. 12, pp. 920–931, Nov. 2005, doi: 10.1038/nmeth815.

2. F. Helmchen and W. Denk, ‘Deep tissue two-photon microscopy’, Nat Methods, vol. 2, no. 12, pp. 932–940, Nov. 2005, doi: 10.1038/nmeth818.

3. J. Mertz, ‘Optical sectioning microscopy with planar or structured illumination’, Nat Methods, vol. 8, no. 10, pp. 811–819, Sep. 2011, doi: 10.1038/nmeth.1709.

4. D. Lim, K. K. Chu, and J. Mertz, ‘Wide-field fluorescence sectioning with hybrid speckle and uniform-illumination microscopy’, Opt Lett, vol. 33, no. 16, p. 1819, Aug. 2008, doi: 10.1364/ol.33.001819.

5. E. H. K. Stelzer et al., ‘Light sheet fluorescence microscopy’, Nature Reviews Methods Primers, vol. 1, no. 1, Nov. 2021, doi: 10.1038/s43586-021-00069-4.

6. C. Dunsby, ‘Optically sectioned imaging by oblique plane microscopy’, Opt Express, vol. 16, no. 25, p. 20306, Nov. 2008, doi: 10.1364/oe.16.020306.

7. G. Hong et al., ‘Through-skull fluorescence imaging of the brain in a new near-infrared window’, Nat Photonics, vol. 8, no. 9, pp. 723–730, Aug. 2014, doi: 10.1038/nphoton.2014.166.

8. V. G. Bandi et al., ‘Targeted multicolor in vivo imaging over 1,000nm enabled by nonamethine cyanines’, Nat Methods, vol. 19, no. 3, pp. 353–358, Feb. 2022, doi: 10.1038/s41592-022-01394-6.

9. O. T. Bruns et al., ‘Next-generation in vivo optical imaging with short-wave infrared quantum dots’, Nat Biomed Eng, vol. 1, no. 4, Apr. 2017, doi: 10.1038/s41551-017-0056.

10. J. A. Carr et al., ‘Shortwave infrared fluorescence imaging with the clinically approved near-infrared dye indocyanine green’, Proceedings of the National Academy of Sciences, vol. 115, no. 17, pp. 4465–4470, Apr. 2018, doi: 10.1073/pnas.1718917115.

11. E. D. Cosco et al., ‘Shortwave infrared polymethine fluorophores matched to excitation lasers enable non-invasive, multicolour in vivo imaging in real time’, Nat Chem, vol. 12, no. 12, pp. 1123–1130, Oct. 2020, doi: 10.1038/s41557-020-00554-5.

12. G. Hong et al., ‘In Vivo Fluorescence Imaging with Ag2S quantum dots in the second near-infrared region’, Angewandte Chemie International Edition, vol. 51, no. 39, pp. 9818–9821, Sep. 2012, doi: 10.1002/anie.201206059.

13. Z. Tao et al., ‘Biological Imaging Using Nanoparticles of Small Organic Molecules with Fluorescence Emission at Wavelengths Longer than 10000.25emnm’, Angewandte Chemie International Edition, vol. 52, no. 49, pp. 13002–13006, Oct. 2013, doi: 10.1002/anie.201307346.

14. K. Welsher et al., ‘A route to brightly fluorescent carbon nanotubes for near-infrared imaging in mice’, Nat Nanotechnol, vol. 4, no. 11, pp. 773–780, Oct. 2009, doi: 10.1038/nnano.2009.294.

15. Q. Yang et al., ‘Rational Design of Molecular Fluorophores for Biological Imaging in the NIR-II Window’, Advanced Materials, vol. 29, no. 12, p. 1605497, Jan. 2017, doi: 10.1002/adma.201605497.

16. F. Wang et al., ‘In vivo non-invasive confocal fluorescence imaging beyond 1,700nm using superconducting nanowire single-photon detectors’, Nat Nanotechnol, vol. 17, no. 6, pp. 653–660, May 2022, doi: 10.1038/s41565-022-01130-3.

17. F. Xia et al., ‘Short-Wave Infrared Confocal Fluorescence Imaging of Deep Mouse Brain with a Superconducting Nanowire Single-Photon Detector’, ACS Photonics, vol. 8, no. 9, pp. 2800–2810, Sep. 2021, doi: 10.1021/acsphotonics.1c01018.

18. Z. Feng et al., ‘Excretable IR-820 for in vivo NIR-II fluorescence cerebrovascular imaging and photothermal therapy of subcutaneous tumor’, Theranostics, vol. 9, no. 19, pp. 5706–5719, 2019, doi: 10.7150/thno.31332.

19. Q. Zhou, Z. Chen, J. Robin, X.-L. Deán-Ben, and D. Razansky, ‘Diffuse optical localization imaging for noninvasive deep brain microangiography in the NIR-II window’, Optica, vol. 8, no. 6, p. 796, Jun. 2021, doi: 10.1364/OPTICA.420378.

20. F. Wang et al., ‘Light-sheet microscopy in the near-infrared II window’, Nat Methods, vol. 16, no. 6, pp. 545–552, May 2019, doi: 10.1038/s41592-019-0398-7.

21. F. Wang et al., ‘In vivo NIR-II structured-illumination light-sheet microscopy’, Proceedings of the National Academy of Sciences, vol. 118, no. 6, Feb. 2021, doi: 10.1073/pnas.2023888118.

22. J. Qi et al., ‘Aggregation-Induced Emission Luminogen with Near-Infrared-II Excitation and Near-Infrared-I Emission for Ultradeep Intravital Two-Photon Microscopy’, ACS Nano, vol. 12, no. 8, pp. 7936–7945, Aug. 2018, doi: 10.1021/acsnano.8b02452.

23. T. Wang et al., ‘Three-photon imaging of mouse brain structure and function through the intact skull’, Nat Methods, vol. 15, no. 10, pp. 789–792, Sep. 2018, doi: 10.1038/s41592-018-0115-y.

24. M. He et al., ‘Aggregation-induced emission nanoprobe assisted ultra-deep through-skull three-photon mouse brain imaging’, Nano Today, vol. 45, p. 101536, Aug. 2022, doi: 10.1016/j.nantod.2022.101536.

25. N. G. Horton et al., ‘In vivo three-photon microscopy of subcortical structures within an intact mouse brain’, Nat Photonics, vol. 7, no. 3, pp. 205–209, Jan. 2013, doi: 10.1038/nphoton.2012.336.

26. H. Wan et al., ‘A bright organic NIR-II nanofluorophore for three-dimensional imaging into biological tissues’, Nat Commun, vol. 9, no. 1, Mar. 2018, doi: 10.1038/s41467-018-03505-4.

27. M. Zhang et al., ‘Bright quantum dots emitting at ∼1,600 nm in the NIR-IIb window for deep tissue fluorescence imaging’, Proceedings of the National Academy of Sciences, vol. 115, no. 26, pp. 6590–6595, Jun. 2018, doi: 10.1073/pnas.1806153115.

28. S. Zhu et al., ‘3D NIR-II Molecular Imaging Distinguishes Targeted Organs with High-Performance NIR-II Bioconjugates’, Advanced Materials, vol. 30, no. 13, p. 1705799, Feb. 2018, doi: 10.1002/adma.201705799.

29. F. Xia, C. Wu, D. Sinefeld, B. Li, Y. Qin, and C. Xu, ‘In vivo label-free confocal imaging of the deep mouse brain with long-wavelength illumination’, Biomed Opt Express, vol. 9, no. 12, p. 6545, Nov. 2018, doi: 10.1364/boe.9.006545.

30. K.-B. Im, S. Han, H. Park, D. Kim, and B.-M. Kim, ‘Simple high-speed confocal line-scanning microscope’, Opt Express, vol. 13, no. 13, p. 5151, 2005, doi: 10.1364/opex.13.005151.

31. R. Engelmann, ‘Faster than real-time: confocal linescan systems provide ideal conditions for millisecond-resolution physiological imaging’, Nat Methods, vol. 3, no. 11, pp. III–V, Oct. 2006, doi: 10.1038/nmeth971.

32. Q. Zhong et al., ‘High-definition imaging using line-illumination modulation microscopy’, Nat Methods, vol. 18, no. 3, pp. 309–315, Mar. 2021, doi: 10.1038/s41592-021-01074-x.

33. E. D. Cosco et al., ‘Bright Chromenylium Polymethine Dyes Enable Fast, Four-Color In Vivo Imaging with Shortwave Infrared Detection’, J Am Chem Soc, vol. 143, no. 18, pp. 6836–6846, May 2021, doi: 10.1021/jacs.0c11599.

34. Y. Zhong et al., ‘In vivo molecular imaging for immunotherapy using ultra-bright near-infrared-IIb rare-earth nanoparticles’, Nat Biotechnol, vol. 37, no. 11, pp. 1322–1331, Nov. 2019, doi: 10.1038/s41587-019-0262-4.

35. Z. Chang et al., ‘Near Infrared-II Fluorescent protein for In-vivo Imaging’, Mar. 2022, doi: 10.1101/2022.03.04.482971.

36. B. D. Hann, E. J. Baldelomar, J. R. Charlton, and K. M. Bennett, ‘Measuring the intrarenal distribution of glomerular volumes from histological sections’, American Journal of Physiology-Renal Physiology, vol. 310, no. 11, pp. F1328–F1336, Jun. 2016, doi: 10.1152/ajprenal.00382.2015.

37. E. Katrukha, ‘Z-stack Depth Color Code’, GitHub repository. GitHub.

38. J. Qi et al., ‘Real-Time and High-Resolution Bioimaging with Bright Aggregation-Induced Emission Dots in Short-Wave Infrared Region’, Advanced Materials, vol. 30, no. 12, p. 1706856, Mar. 2018, doi: 10.1002/adma.201706856.

39. J. Meng et al., ‘Mapping physiological and pathological functions of cortical vasculature through aggregation-induced emission nanoprobes assisted quantitative, in vivo NIR-II imaging’, Biomaterials Advances, vol. 136, p. 212760, May 2022, doi: 10.1016/j.bioadv.2022.212760.

40. B. A. Molitoris and R. M. Sandoval, ‘Intravital multiphoton microscopy of dynamic renal processes’, American Journal of Physiology-Renal Physiology, vol. 288, no. 6, pp. F1084–F1089, Jun. 2005, doi: 10.1152/ajprenal.00473.2004.

41. L. Wei, Z. Chen, and W. Min, ‘Stimulated emission reduced fluorescence microscopy: a concept for extending the fundamental depth limit of two-photon fluorescence imaging’, Biomed Opt Express, vol. 3, no. 6, p. 1465, May 2012, doi: 10.1364/boe.3.001465.

42. A. G. York et al., ‘Instant super-resolution imaging in live cells and embryos via analog image processing’, Nat Methods, vol. 10, no. 11, pp. 1122–1126, Oct. 2013, doi: 10.1038/nmeth.2687.

